# Minmers are a generalization of minimizers that enable unbiased local Jaccard estimation

**DOI:** 10.1101/2023.05.16.540882

**Authors:** Bryce Kille, Erik Garrison, Todd J Treangen, Adam M Phillippy

## Abstract

1

**Motivation:** The Jaccard similarity on *k*-mer sets has shown to be a convenient proxy for sequence identity. By avoiding expensive base-level alignments and comparing reduced sequence representations, tools such as MashMap can scale to massive numbers of pairwise comparisons while still providing useful similarity estimates. However, due to their reliance on minimizer winnowing, previous versions of MashMap were shown to be biased and inconsistent estimators of Jaccard similarity. This directly impacts downstream tools that rely on the accuracy of these estimates.

**Results:** To address this, we propose the *minmer* winnowing scheme, which generalizes the minimizer scheme by use of a rolling minhash with multiple sampled *k*-mers per window. We show both theoretically and empirically that minmers yield an unbiased estimator of local Jaccard similarity, and we implement this scheme in an updated version of MashMap. The minmer-based implementation is over 10 times faster than the minimizer-based version under the default ANI threshold, making it well-suited for large-scale comparative genomics applications.

**Availability:** MashMap3 is available at https://github.com/marbl/MashMap

## 2 Introduction

The recent deluge of genomic data accelerated by population-scale long-read sequencing efforts has driven an urgent need for scalable long-read mapping and comparative genomics algorithms. The completion of the first Telomere-to-Telemore (T2T) human genome Nurk *et al*. (2022) and the launch of the Human Pangenome Project Wang *et al*. (2022a) have paved the way to mapping genomic diversity at unprecedented scale and resolution. A key goal when comparing a newly sequenced human genome to a reference genome or pangenome is to accurately identify homologous sequences, that is, DNA sequences that share a common evolutionary source.

Algorithms for pairwise sequence alignment, which aim to accurately identify homologous regions between two sequences, have continued to advance in recent years Marco-Sola *et al*. (2021). While a powerful and ubiquitous computational tool in computational biology, exact alignment algorithms are typically reserved for situations where the boundaries of homology are known *a priori*, due to their quadratic runtime costs and inability to model nonlinear sequence relationships such as inversions, translocations, and copy number variants. Because of this, long-read mapping or whole-genome alignment methods must first identify homologous regions across billions of nucleotides, after which the exact methods can be deployed to compute a base-level “gapped” read alignment for each region. To efficiently identify candidate mappings, the prevailing strategy is to first sample *k*-mers and then identify consecutive *k*-mers that appear in the same order for both sequences: known as “seeding” and “chaining”, respectively.

For many use cases, an exact gapped alignment is not needed and only an estimate of sequence identity is required. As a result, methods have been developed which can predict sequence identity without the cost of computing a gapped alignment. Jaccard similarity, a metric used for comparing the similarity of two sets, has found widespread use for this task, especially when combined with locality sensitive hashing of *k*-mer sets Ondov *et al*. (2016); Brown and Irber (2016); Ondov *et al*. (2019); Jain *et al*. (2017, 2018a); Baker and Langmead (2019); Shaw and Yu (2023). By comparing only *k*-mers, the Jaccard can be used to estimate the average nucleotide identity (ANI) of two sequences without the need for an exact alignment Ondov *et al*. (2016, 2019); Blanca *et al*. (2022).

To accelerate mapping and alignment, *k*-mers from the input sequences are often down-sampled using a “winnowing scheme” in a way that reduces the input size while still enabling meaningful comparisons. For example, both MashMap Jain *et al*. (2017, 2018a) and Minimap Li (2018) use a minimizer scheme Roberts *et al*. (2004), which selects only the *smallest k*-mer from all *w*-length substrings of the genome. Of relevance to this study, MashMap2 then uses these minimizers to approximate the Jaccard similarity between the mapped sequences, and these estimates have been successfully used by downstream methods such as FastANI Jain *et al*. (2018b) and MetaMaps Dilthey *et al*. (2019).

However, a recent investigation noted limitations of the “winnowed minhash” scheme introduced by MashMap Belbasi *et al*. (2022). Although the original MashMap paper notes a small, but negligible bias in its estimates Jain *et al*. (2017), Belbasi *et al*. proved that no matter the length of the sequences, the bias of the minimizer-based winnowed minhash estimator is never zero Belbasi *et al*. (2022).

To address this limitation, we propose a novel winnowing scheme, the “minmer” scheme, which is a generalization of minimizers that allows for the selection of multiple *k*-mers per window. We define this scheme, characterize its properties, and provide an implementation in MashMap3. Importantly, we show that minmers, unlike minimizers, enable an unbiased prediction of the local Jaccard similarity.

## 3 Preliminaries

Let Σ be an alphabet and *𝒮*_*k*_(*S*) : Σ^+^ → {Σ^*k*^} ^+^ be a function which maps a sequence *S* to the set of all *k*-mers in *S*. Similarly, given a sequence *S*, we define 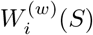 as the sequence of *w k*-mers in *S* starting at the *i*th *k*-mer. When *w* and *S* are clear from context, we use *W*_*i*_. We use the terms sequence and string interchangeably.

### 3.1 Jaccard similarity and the minhash approximation

Given two sets *A* and *B*, their Jaccard similarity is defined as 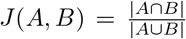. The Jaccard similarity between two sequences *R* and *Q* can be computed as *J* (𝒮_*k*_(*R*), 𝒮_*k*_(*Q*)) for some *k*-mer size *k*.

However, computing the exact Jaccard for 𝒮_*k*_(*R*) and 𝒮_*k*_(*Q*) is not an efficient method for determining similarity for long reads and whole genomes. Instead, the minhash algorithm provides an estimator for the Jaccard similarity while only needing to compare a fraction of the two sets. Assuming *U* is the universe of all possible elements and *π* : *U* → | *U* | is a function which imposes a randomized total order on the universe of elements, we have that

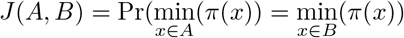

This equivalency, proven by Broder (1997), is key to the minhash algorithm and yields an unbiased and consistent Jaccard estimator *Ĵ* with the help of a sketching function *π*_*s*_. Let *π*_*s*_ return the lowest *s* items from the input set according to the random total order *π*. Then we define the minhash as

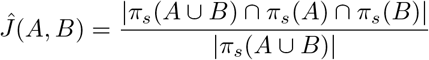

Importantly, this Jaccard estimator has an expected error that scales with 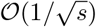 and is therefore independent of the size of the original input sets. While there are a number of variants of minhash which provide the same guarantee Cohen (2016), we will be using the “bottom-*s* sketch” (as opposed to the *s*-mins and *s*-partition sketch) since it ensures a consistent sketch size regardless of the parameters and requires only a single hash computation per element of *𝒮*_*k*_. Additionally, the simplicity of the bottom-*s* sketch leads to a streamlined application of the sliding window model, which we describe next.

### 3.2 Winnowing

While sequences can be reduced into their corresponding sketch via the method described above, this is a *global* sketch and it is difficult to determine *where* two sequences share similarity. In order to perform local mapping, Schleimer *et al*. (2003) and Roberts *et al*. (2004) independently introduced the concept of *winnowing* and *minimizers*. In short, given some total ordering on the *k*-mers, a window of length *w* is slid over the sequence and the element with the lowest rank in each window (the *minimizer*) is selected, using the left-most position to break ties Roberts *et al*. (2004). By definition, winnowing ensures that at least one element is sampled per window and therefore there is never a gap of more than *w* elements between sampled positions. Here, we extend the winnowing concept to allow the selection of more than one element per window (the *minmers*), and we refer to the set of all minmers and/or their positions as the *winnowed* sequence.

#### 3.2.1 Winnowing scheme characteristics

##### Definition 3.1

*A winnowing scheme has a* (*w, s*)*-window guarantee if for every window of w k-mers, there are at least* max(#_*distinct*_, *s*) *k-mers sampled from the window, where* #_*distinct*_ *is the number of distinct k-mers in the window*.

This definition is more general than the commonly used *w*-window guarantee, which is equivalent to the (*w*, 1)-window guarantee. While not all winnowing schemes must have such a guarantee, this ensures that no area of the sequence is under-sampled. Shaw and Yu (2022) recently provided an analytical framework for winnowing schemes and showed that mapping sensitivity is related to the distribution of distances (or *spread*) between sampled positions, and precision is related to the proportion of unique values relative to the total number of sampled positions. As the over-arching goal of winnowing is to reduce the size of the input while preserving as much information as possible, winnowing schemes typically aim to optimize the precision/sensitivity metrics given a particular density.

##### Definition 3.2

*The density d of a winnowing scheme is defined as the expected frequency of sampled positions from a long random string, and the density factor d*_*f*_ *is defined as the expected number of sampled positions in a window of w* + 1 *k-mers*.

There has been significant work on improving the performance of minimizers by identifying orderings that reduce the density factor Marçais *et al*. (2017). Minimizer schemes which use a uniformly random ordering have a density factor of *d*_*f*_ = 2 and recent schemes like Miniception Zheng *et al*. (2020) and PASHA Ekim *et al*. (2020) are able to obtain density factors as low as 1.7 for certain values of *w* and *k*.

For the remainder of this work, we will assume that *w* ≪ 4^*k*^, i.e. the windows are not so large that we expect duplicate *k*-mers in a random string. This ensures that each *k*-mer in a window has probability *s/w* of being in the sketch for that window.

#### 3.2.2 Winnowing scheme hierarchies

Recent winnowing methods have focused on schemes that select at most a single position per window, which simplifies analyses but restricts the universe of possible schemes. Minimizers belong to the class of *forward* winnowing schemes, where the sequence of positions sampled from adjacent sliding windows is non-decreasing Marçais *et al*. (2018). More general is the concept of a *w*-local scheme Shaw and Yu (2022), defined on windows of *w* consecutive *k*-mers but without the forward requirement. Non-forward schemes are more powerful and are not limited by the same density factor bounds as forward schemes. While the need of non-forward schemes to “jump back” in order to obtain lower sampling densities is acknowledged by Marçais *et al*. (2018), there are currently no well-studied, non-forward, *w*-local schemes.

### 3.3 MashMap

MashMap is a minimizer-based tool for long-read and whole-genome sequence homology mapping that is designed to identify all pairwise regions above some sequence similarity cutoff Jain *et al*. (2017, 2018a). Specifically, for a reference sequence *R* and a query sequence *Q* comprised of *w k*-mers, MashMap aims to find all positions *i* in the reference such that *J* (*A, B*_*i*_) ≥ *c*, where *A* = *𝒮*_*k*_(*Q*) and 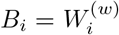, and *c* is the sequence similarity cutoff. For ease of notation, we will use *B* to refer to the sequence of *k*-mers from the reference sequence *R*. Importantly, MashMap only requires users to specify a minimum segment length and minimum sequence identity threshold, and the algorithm will automatically determine the parameters needed to return all mappings that meet this criteria with parameterized confidence under a binomial mutation model.

Here we replace the minimizer-based approach of prior versions of MashMap with minmers. While the problem formulation remains the same, our method for computing the reference index and filtering candidate mappings is novel. We will first introduce the concept of minmers, which enable winnowing the input sequences while still maintaining the *k*-mers necessary to compute an unbiased Jaccard estimation between any two windows of length at least *w*. We will then discuss the construction of the reference index and show how query sequences can be efficiently mapped to the reference such that their expected ANI is above the desired threshold.

## 4 The minmer winnowing scheme

Minmers are a generalization of minimizers that allow for the selection of more than one minimum value per window. The relationship between minmers and minimizers was noted by Berlin *et al*. (2015) but as a global sketch and without the use of a sliding window. Here we formalize a definition of the minmer winnowing scheme.

### Definition 4.1

*Given a tuple* (*w, s, k, π*), *where w, k and s are integers and π is an ordering on the set of all k-mers, a k-mer in a sequence is a minmer if it is one of the smallest s k-mers in any of the subsuming windows of w k-mers*.

Similar to other *w*-local winnowing schemes, ties between *k*-mers are broken by giving priority to the leftmost *k*-mer. From the definition, it follows that by letting *s* = 1 we obtain the definition of the minimizer scheme. Compared to minimizers with the same *w* value, minmers guarantee that at least *s k*-mers will be sampled from each window. However, as a non-forward scheme, a minmer may be one of the smallest *s k*-mers in two non-adjacent windows, yet not one of the smallest *s k*-mers in an intervening window (Figure 1). To account for this and simplify development of this scheme, we define a *minmer interval* to be the interval for which the *k*-mer at position *i* is a minmer for all windows starting within that interval. Thus, a single *k*-mer may have multiple minmer intervals starting at different positions.

**Figure 1:**
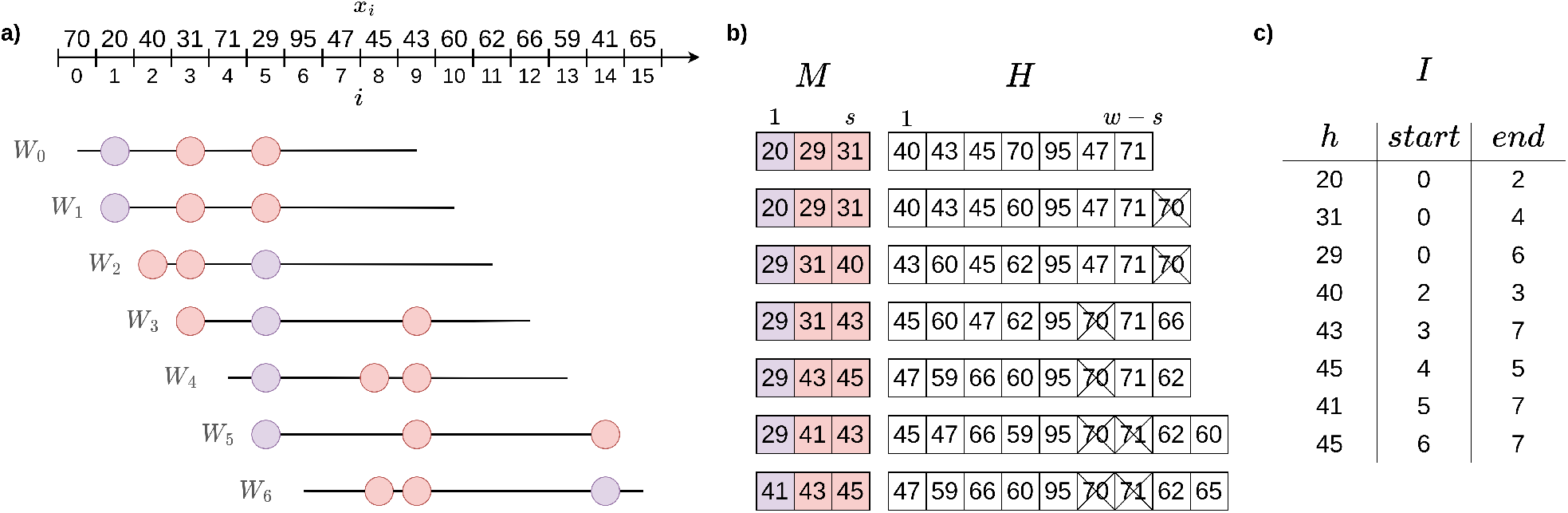
Constructing the rolling minhash index. **(a)** A sliding window *Bi* of length *w* = 10 is moved over the hashes of all *k*-mers. At each position *i* of the sliding window, the positions with the *s* = 3 lowest hash values are marked as minmers. The 3 minmers for each window are highlighted with colored circles, with the smallest hash in each window (the minimizer) highlighted in purple. **(b)** The values of the hashes in the map *M* and heap *H* as the window slides over the sequence. The expired *k*-mers in the heap are crossed out. **(c)** The final sorted minmer interval index *I*.

### Definition 4.2

*A tuple* (*i, a, b*) *is a minmer interval for a sequence S if the k-mer at position i is a minmer for all windows W*_*j*_ *where j* ∈ [*a, b*), *but not W*_*a−*1_ *or W*_*b*_.

Any window *W*_*j*_ may contain more than *s* minmers, and so to naively compute the Jaccard between a query and *W*_*j*_ would require identification of the *s* smallest *k*-mers in *W*_*j*_. Minmer intervals are convenient because for any window start position *j*, the *s* smallest *k*-mers in *W*_*j*_ are simply the ones whose minmer intervals contain *j*. Thus, indexing *S* with minmer intervals enables the efficient retrieval of the smallest *s k*-mers for any window without additional sorting or comparisons.

Another benefit of minmer intervals is that the smallest *s k*-mers for any window of length *w*′ > *w* are guaranteed to be a subset of the combined (*w, s*)-minmers contained in that window. This subset can be easily computed with minmer intervals, since the set of (*w, s*)-minmer intervals that overlap with the range [*i, i* + *w*′ − *w*] are also guaranteed to include the *s* smallest *k*-mers of the larger window, and the overlapping minmer intervals can be inspected to quickly identify them.

### 4.1 Constructing the rolling minhash index

In this section, we will describe our rolling bottom-*s* sketch algorithm for collecting minmers and their corresponding minmer intervals. Popic and Batzoglou (2017) proposed a related rolling minhash method for short-read mapping, but using an *s*-mins scheme without minmer intervals. For the remainder of the section, we will assume no duplicate *k*-mers in a window and an ideal uniform hash function which maps to [0, 1]. Duplicate *k*-mers are handled in practice by keeping a counter of the number of active positions for a particular *k*-mer, similar to the original MashMap implementation Jain *et al*. (2017). Minmer intervals longer than the window length sometimes arise due to duplicate *k*-mers and are split into adjacent windows of length at most *w*. This bound on the minmer interval length is necessary for the mapping step.

For ease of notation, we now consider *B* as a sequence of *k*-mer hash values *x*_0_, *x*_1_, …, *x*_*n*_ where each *x*_*i*_ ∈ [0, 1] and refer to these elements as hashes and *k*-mers interchangeably. We use a min-heap *H* and a sorted map *M*, both ordered on the hash values, to keep track of the rolling minhash index. As the window slides across *B, M* will contain the minmer intervals for the lowest *s* hashes in the window and *H* will contain the remaining hashes in the window. We denote the minmer interval of a hash *x* in *M* by *M* [*x*]^(*start*)^ and *M* [*x*]^(*end*)^. In practice, *H* may contain “expired” *k*-mers which are no longer part of the current window, however by storing the *k*-mer position as well, we can immediately discard such *k*-mers whenever they appear at the top of the heap. To prevent expired *k*-mers from accumulating, all expired *k*-mers from the heap are pruned whenever the heap size exceeds 2*w*. After initialization of *H* and *M* with the first *w k*-mers of *B*, we begin sliding the window for each consecutive position *i* and collect the minmer intervals in an index *I*. For each window *B*_*i*_, there will be a single “exiting” *k*-mer *x*_*i−*1_ and a single “entering” *k*-mer *x*_*i*+*w−*1_, each of which may or may not belong to the lowest *s k*-mers. Therefore, we have four possibilities, examples of which can be seen in Figure 1.

1. *x*_*i*−1_ > max(*M*) and *x*_*i*+*w*−1_ > max(*M*) Neither the exiting nor entering *k*-mer is in the sketch. Insert *x*_*i*+*w*−1_ into *H*.
2. *x*_*i*−1_ > max(*M*) and *x*_*i*+*w*−1_ < max(*M*) The exiting *k*-mer was not in the sketch, but the entering *k*-mer will be. Since the incoming *k*-mer *x*_*i*+*w*−1_ enters the sketch, the largest element in the sketch must be removed. Therefore, *M* [max(*M*)]^(*end*)^ is set to *i* and the the minmer interval is appended to the index *I*. max(*M*) is then removed from *M* and the new *k*-mer *x*_*i*+*w*−1_ is inserted to *M*, marking *M* [*x*_*i*+*w*−1_]^(*start*)^ = *i*.
3. *x*_*i*−1_ ≤ max(*M*) and *x*_*i*+*w*−1_ > max(*M*) The exiting *k*-mer was in the sketch, but the entering *k*-mer will not be. Since the exiting *k*-mer *x*_*i*−1_ was a member of the sketch, set *M* [*x*_*i*−1_]^(*end*)^ = *i*, remove *M* [*x*_*i*−1_] from *M* and append it to *I*, and insert *x*_*i*+*w*−1_ into *H*. At this point, *M* = *s* 1, as we removed an element from the sketch but did not replace it. To fill the empty sketch position, *k*-mers are popped from *H* until a *k*-mer *x* which has not expired is obtained. This *k*-mer is added to *M*, setting *M* [*x*]^(*start*)^ = *i*.
4. *x*_*i*−1_ ≤ max(*M*) and *x*_*i*+*w*−1_ ≤ max(*M*) Both the exiting and entering *k*-mers are in the sketch. As before, set *M* [*x*_*i*−1_]^(*end*)^ = *i* and remove *M* [*x*_*i*−1_] from *M* and append it to *I*. The entering *k*-mer belongs in the sketch, so set *M* [*x*_*i*+*w*−1_]^(*start*)^ = *i*.

Our implementation of *M* uses a balanced binary tree and *H* is pruned in 𝒪 (*w*) time at most every *w k*-mers and therefore the amortized time complexity of each sliding window update is 𝒪 (log(*w*)). In order to efficiently use the index for mapping, we sort *I* based on the start positions of the minmers. In addition to *I*, we compute a reverse lookup table *T* which maps hash values to ordered lists of start and end points of minmer intervals for that hash value. Overall, the indexing time requires 𝒪 (*n* log(*w*) + |*I*| log(|*I*|)), where | I | is estimated to be 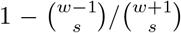, as shown in section 5.1.2.

### 4.2 Querying the rolling minhash index

MashMap computes mappings in a two-stage process. In the first stage, all regions within the reference that may contain a mapping satisfying the desired ANI constraints are obtained. In the second stage, the minhash algorithm is used to estimate the Jaccard for each candidate mapping position *i* produced by the first stage. As the second stage is the most computationally intensive step, we introduce both a new candidate region filter and a more efficient minhash computation to improve overall runtime. We assume here that query sequences are *w k*-mers long. In practice, sequences longer than *w* are split into windows of *w k*-mers, mapped independently, and then chained and filtered as described in Jain *et al*. (2018a).

#### 4.2.1 Stage 1: Candidate region filter

First, the query sequence *A* is winnowed using a min-heap to obtain the *s* lowest hash values. All *m* minmer intervals in the reference with matching hashes are obtained from *T* and a sorted list *L* is created in 𝒪 (*m* log(*s*)) time, where *L* consists of all minmer start and end positions. In this way, we can iterate through the list and keep a running count of the overlapping minmer intervals by incrementing the count for each start-point and decrementing the count for each end-point.

Unlike the previous versions of MashMap that look for all mappings above a certain ANI threshold, MashMap3 provides the option to instead filter out all mappings which are not likely to be within Δ_*ANI*_ of the best predicted mapping ANI. This significantly reduces the number and size of the candidate regions passed on to the more expensive second stage.

Let *Y*_*i*_ be a random variable representing the numerator of the minhash formula for *A* and *B*_*i*_. Given *c*_*i*_ = |*π*_*s*_(*A*) ⋂ *π*_*s*_(*B*_*i*_) |, we observe that *Y*_*i*_ is distributed hypergeometrically, where we have *s* success states in a population of 2*s* − *c*_*i*_ states (proof in Supplementary Materials). Let *z* = arg max_*i*_ *c*_*i*_ be a position with the maximum intersection size over all *B*_*i*_, i.e. the position in *B* that overlaps with the most selected minmer intervals. We can now find a minimum intersection size *τ* such that for any *c*_*i*_ < τ,

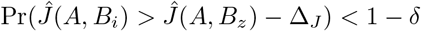

where Δ_*J*_ is the difference in the Jaccard that corresponds to an ANI value Δ_ANI_ less than the ANI value predicted by *Ĵ* (*A,Bz*) and *δ* is a desired confidence level. To calculate this probability, we can use the following summation

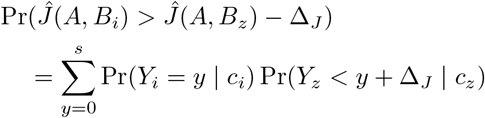

For each intersection size, we can identify a cutoff in 𝒪 (*s* log(*s*)) time. As a preprocessing step, we compute cutoffs for each of the *s* possible intersection sizes at the indexing stage. Candidate regions that are unlikely to have an ANI within Δ_*ANI*_ of the best predicted ANI are then pruned. The default Δ_*ANI*_ and *δ* confidence parameters of MashMap3 are 0 and 0.999, respectively, as in many cases the lower scoring mappings for a segment are filtered out by the plane-sweep filtering method of MashMap described in Jain *et al*. (2018a).

We compute two passes over the interval endpoints in *L*. In the first pass of stage 1, the maximum intersection size *c*_*z*_ is obtained. In the second pass, candidate mappings whose intersection is above the cutoff derived from *c*_*z*_ are obtained. Consecutive candidate mappings are grouped into candidate regions and passed to stage 2.

#### 4.2.2 Stage 2: Efficiently computing the rolling minhash

Given a candidate region [*a, z*), the goal of stage 2 is to calculate the minhash for all *A, B*_*i*_ pairs for *i* ∈ [*a, z*). In order to track the minhash of *A* and *B*_*i*_ for each *i*, MashMap2 previously used a sorted map to track all active seeds in each window. We improve upon this by observing that the minhash can be efficiently tracked using only *π*_*s*_(*A*), *π*_*s*_(*A*) ∩ *π*_*s*_(*B*_*i*_), and the number of minmers from *π*_*s*_(*B*_*i*_) in-between each consecutive pair of minmers from *π*_*s*_(*A*). To do so, MashMap3 uses an array *V* = (−1, 0, 0), (*x*_1_, *α*_1_, *β*_1_), (*x*_2_, *α*_2_, *β*_2_), …, (*x*_*s*_, *α*_*s*_, *β*_*s*_) where each *x*_*j*_ represents one of the *s* minmer hash values from *π*_*s*_(*A*) in increasing order and for each *i* ∈ [*a, z*), the values *α*_*j*_ and *β*_*j*_ are

- *α*_*j*_ = 1 if *x*_*j*_ ∈ π_*s*_(*B*_*i*_) else 0
- *β*_*j*_ = 1 + |{x ∈ π_*s*_(*B*_*i*_) s.t. *x*_*j*−1_ < *x* < *x*_*j*_}|

We can imagine *V* as a set of *s* buckets labeled by the *s* corresponding hash values of *A* and sorted in increasing order. At each position *i* ∈ [*a, z*), each bucket *j* holds *x*_*j*_ and all *β*_*j*_ − 1 reference minmers in *π*_*s*_(*B*_*i*_), which are between *x*_*j*_ and *x*_*j*−1_. A bucket is marked “good” (*α*_*j*_ → 1) if *x*_*j*_ ∈ *π*_*s*_(*B*_*i*_). It remains to find the largest integer *p*_*i*_ such that the number of minmers in the first *p*_*i*_ buckets is at most *s*. Given *p*_*i*_, the numerator of the minhash formula, *Y*_*i*_, is the number of “good” buckets in the first *p*_*i*_ buckets.

For a candidate region [*a, z*), we initialize *V* by inserting all of the minmers from the reference index whose intervals overlap with *a* and set

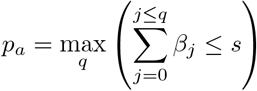

It follows that 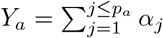

In order to keep track of intervals which overlap with the current position, we use a min-heap *H* sorted on interval endpoints. We then continue to iterate through minmer intervals from the reference in order based on their start points, stopping once the intervals no longer overlap with [*a, z*). For each minmer interval starting at *i* ∈ [*a* + 1, *z*), we pop intervals from *H* that end at or before *i*. For each interval popped from *H*, we update *V* in 𝒪 (log(*s*)) time through a binary search, decrementing the corresponding *β*_*j*_ and setting *α*_*j*_ = 0 if the interval represents a shared minmer. The new interval is added in a similar manner and the necessary *α* and *β* values are updated. After *V* is updated, *p*_*i*_ is updated from *p*_*i*−1_ by incrementing or decrementing until it is the maximal value such that 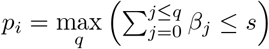. By keeping track of *p*_*i*−1_ and the sums 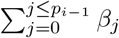 and 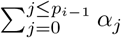, the new *p*_*i*_ and corresponding sums are updated in constant time per window.

While the MashMap3 implementation of the second filtering stage still requires 𝒪 (log(*s*)) time to update the minhash for each sliding window within the candidate region, it is significantly more efficient than MashMap2’s ordered map in practice due to *V* being a static data structure in contiguous memory, only requiring updates to counters.

#### 4.2.3 Early termination of stage 2

Instead of computing the stage 2 step for each candidate region obtained in the first stage, we aim to terminate the second stage once we have confidently identified all mappings whose predicted ANI is within Δ_ANI_ of the best predicted ANI. We do this by sorting the candidate regions in decreasing order of their maximum interval overlap size obtained in stage 1. The stage 2 minhash calculation is then performed on each candidate region in order, keeping track of the best predicted ANI value seen. Let *κ* be numerator of the minhash that corresponds to an ANI value Δ_ANI_ less than the best predicted ANI value seen so far. Then, given a candidate region with a maximum overlap size of *c*_*i*_ *κ*, we know that Pr(*Y*_*i*_ ≥ *κ*) = 0 and therefore no more candidate regions can contain mappings whose predicted ANI is within Δ_ANI_ of the predicted ANI of the best mapping.

## 5 Results

### 5.1 Characteristics of the minmer scheme

Here we provide formulas for the density of minmers and minmer intervals and an approximation for the distance between adjacent minmers. Proofs of the formulas are presented in the Supplementary Materials. We then compare these formulas to results on both simulated and empirical sequences. For the simulated dataset, we generated a sequence of 1 million uniform random hash values. For the empirical dataset, we used MurmurHash to hash the sequence of *k*-mers in the recently-completed human Y-chromosome Rhie *et al*. (2022) with *k* = 18.

#### 5.1.1 Minmer density

To obtain the formula for the minmer density, we consider how the rank of a random *k*-mer changes with each consecutive window that contains it. As a result, we have a distribution of the rank of a random *k*-mer throughout consecutive sliding windows. This distribution enables us to not only obtain the density (Figure 2), but also determine other characteristics such as the likelihood of being a minmer given some initial rank *r*_1_ or given a hash value *z*.

**Figure 2:**
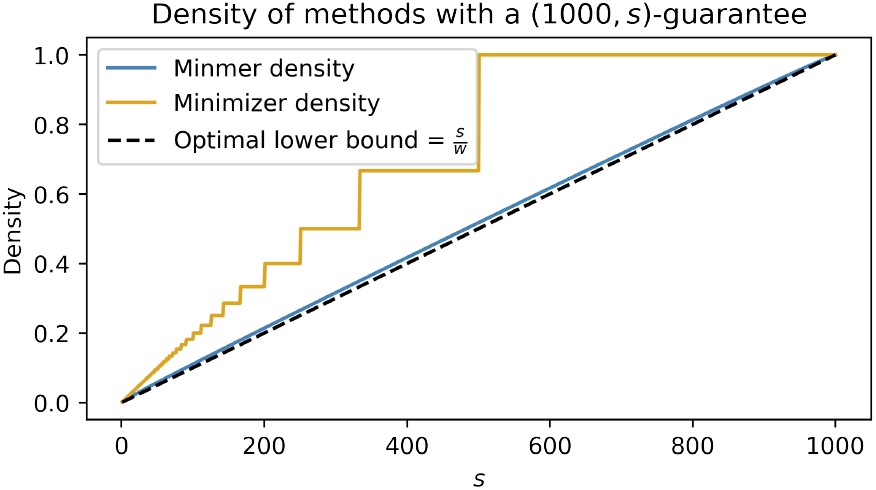
The density of a (1000, *s*)-minmer scheme compared to a *w*′-minimizer scheme which also yields a (1000, *s*)-window guarantee. To ensure that the minimizer scheme satisfies the (1000, *s*) window guarantee, the minimizer scheme is set with *w*′ = [1000/*s*].

**Theorem 5.1**

*Let d*_(*w,s*)_ *be the expected density of* (*w, s*)*-minmers in a random sequence. Then*,

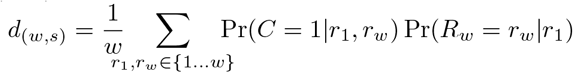

*where R*_*w*_|*r*_1_ ∼ *BetaBinomial*(*r*_1_, *w* − *r*_1_ + 1) *and*

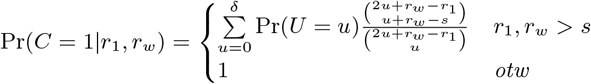

*where U* ∼ *Hypergeometric*(*w* − 1, *r*_1_ − 1, *w* − *r*_*w*_) *and δ* = min(*r*_1_ − 1, *w* − *r*_*w*_).

#### 5.1.2 Minmer interval density

**Theorem 5.2**

*Let* 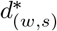 *be the density of* (*w, s*)*-minmer intervals in a random sequence. Then*,

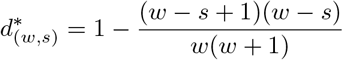

As expected, letting *s* = 1 yields the same density as minimizers, 2*/*(*w* + 1), and a similar formula appears when determining the probability of observing *s* consecutive unsampled *k*-mers under the the minimizer scheme Spouge (2022). As the number of minmers is a strict lower bound on the number of minmer intervals, this result also gives an upper bound on the density of (*w, s*)-minmers.

#### 5.1.3 Minmer window guarantee

As the main difference between minimizers and minmers is the window guarantee, it is important to observe the difference in the density of the minmer scheme compared to a minimizer scheme which also satisfies the (*w, s*)-window guarantee. In Figure 2, we consider the case where we have a (1000, *s*)-minmer scheme and a *w*′-minimizer scheme, where *w*′ is set to obtain the same window guarantee of the minmer scheme by letting *w*′ = [1000/*s* ]. We observe that for sketch sizes other than 1 and 1000, for which the density of the schemes are equal, the density of the minmer scheme is strictly less than the density of the corresponding minimizer scheme. For some values of *s*, the density of the [1000/*s*] -minimizer scheme is over 70% larger than the (1000, *s*)-minmer scheme.

#### 5.1.4 Minmer spread

Let *G*_*i*_ be the distance between the *i*^th^ selected minmer and the (*i* + 1)^th^ selected minmer. For a (*w, s*)-minmer scheme with a density factor *d*_*f*_, we have that

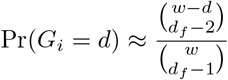

To see how well this approximation holds, we plot the results on both empirical and simulated data in Supplemental Figure 2.

### 5.2 ANI prediction ideal sequences

We replicated the experiments for Table 1 of Belbasi *et al*. (2022) using the minmer-based MashMap3 (commit 0b47608), with the exception that we report the mean predicted sequence divergence as opposed to the median. For each divergence rate *r* ∈ {0.01, 0.05, 0.10}, 100 random windows of 10,000 base pairs were selected from the *Escherichia coli* genome and 10, 000*r* positions were selected at random and mutated, ensuring that no duplicate *k*-mers were generated. The reads were mapped back to the reference *E. coli* genome and the predicted divergence was compared to the ground truth (Figure 3).

**Table 1:** Metrics for simulated Nanopore read mapping to the human genome. Minmer and minimizer-based MashMap implementations as well as Minimap2 were used to map simulated reads from the human reference genome using Pbsim Ono *et al*. (2013).

| Dataset | Minimap2 |  |  |  | MashMap2 |  |  |  | MashMap3 |  |  |  |
| --- | --- | --- | --- | --- | --- | --- | --- | --- | --- | --- | --- | --- |
|  | CPU time (m) | Memory (Gb) | ME | MAE | CPU time (m) | Memory (Gb) | ME | MAE | CPU time (m) | Memory (Gb) | ME | MAE |
| ONT-99 | 154.20 | **9.89** | -0.25 | 0.34 | 80.27 | 9.92 | -0.27 | 0.29 | **33.64** | 13.07 | **0.03** | **0.17** |
| ONT-98 | 147.29 | **9.89** | -0.36 | 0.52 | 82.46 | 9.92 | -0.33 | 0.39 | **35.13** | 13.09 | **0.06** | **0.29** |
| ONT-95 | 96.35 | **9.89** | -0.46 | 0.81 | 106.81 | 9.92 | -0.25 | **0.59** | **42.81** | 13.10 | **0.21** | 0.62 |

**Figure 3:**
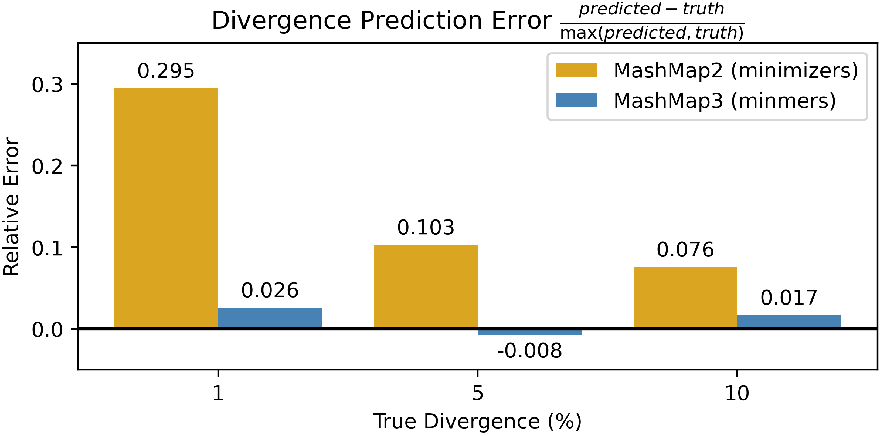
Eliminating the bias in MashMap. The experiments from Table 1 of Belbasi *et al*. (2022) were replicated. Divergence, defined as 1-ANI, was predicted across 100 sequences for both MashMap2 and MashMap3 using a density of 0.009 (*L* = 10, 000, *s* = 78). In the case of identifying sequence divergence of very closely related genomes, the minmer scheme reduces relative prediction error from 29% to 2%

The parameters of the minmer-based MashMap3 were set to obtain a similar numbers of sampled *k*-mers as the minimizer-based MashMap2 under MashMap2’s default settings, resulting in a density of 0.009 for both tools. As expected, the results show that the ANI values predicted by the minmer scheme are significantly closer to the ground truth than those predicted by the minimizer scheme. Notably, in the case where the true divergence was 1%, the relative error is reduced from 29% to 2% (Figure 3).

### 5.3 ANI prediction on simulated reads

In addition to the ANI prediction measurements from Belbasi *et al*. (2022), we also simulated reads from the human T2T-CHM13 reference genome Nurk *et al*. (2022) at varying error rates to determine the accuracy of the ANI predictions. We compared the minmer-based MashMap3 against the minimizer-based MashMap2 with similar densities for each run as well as against Minimap2 Li (2018). Minimap2 was run in its default mode with -x map-ont set which, like MashMap, computes approximate mappings and estimates the alignment identity. MashMap2 was modified to use the binomial model for estimating the ANI from the Jaccard estimator which has been shown to be more accurate Belbasi *et al*. (2022).

We used Pbsim Ono *et al*. (2013) to simulate three datasets: “ONT-95”, “ONT-98”, and “ONT-99”, where the number following the dash represents the average ANI across reads. The standard deviation of the error rates was set to 0, and the ratio of matches, insertions, and deletions was set to 20:40:40, respectively, to ensure that mapped regions would, on average, be the same length as the reads. For each dataset, 5,000bp reads were generated with the CLR profile at a depth of 2, resulting in 1.25 million reads for each dataset. The mappings output by the different methods were parsed and the predicted ANI was compared to the gap-compressed ANI of the ground-truth mapping. The results of the simulations can be seen in Table 1.

For MashMap2 and MashMap3, we used a *k*-mer size of 19 and set the MashMap2 minimizer *w* to 89 and minmer *s* to *s* = 100 to obtain a density of 0.0222 for both tools. The ANI cutoff was set to 94%, 93%, and 90% for the ONT-99, ONT-98, and ONT-95 datasets, respectively. The indexing times for Minimap2, MashMap2, and MashMap3 were 1.7, 2.8, and 9.8 minutes, respectively.

### 5.4 ANI prediction on mammalian genome alignments

To test the performance of MashMap3 at the genome-mapping scale, we computed mappings between the T2T human reference genome and reference genomes for chimpanzee Kronenberg *et al*. (2018) and macaque Warren *et al*. (2020). In absence of ground truth ANI values, we used wfmash Guarracino *et al*. (2021) to compute the gap-compressed ANI of the segment mappings output by MashMap and report the results of the mappings with ≥ 80% complexity in Table 2. For a small proportion of segment mappings output by MashMap2 and MashMap3, wfmash did not produce an alignment. When the ANI threshold is 85%, these cases accounted for 0.07% of chimpanzee mappings and 0.3% macaque mappings. When the ANI threshold was 90% or 95%, less than 0.01% of mappings were not aligned with wfmash for both chimpanzee and macaque. We consider these mappings as false positives. For the ANI thresholds of 95%, 90%, and 85%, the winnowing scheme densities were set to 0.043, 0.053, and 0.064, respectively.

**Table 2:** Comparison of MashMap2 and MashMap3 for identifying mappings between pairs of mammalian genomes. MashMap2 and MashMap3 were used to align the human reference genome to chimpanzee and macaque genomes. The ME and MAE metrics shown are for query segments with at least 80% *k*-mer complexity. Corresponding metrics for low-complexity mappings can be found in Supplementary Table 1. ^*^Sampling bias leads to ANI over-estimation. See discussion for details.

| Query Species | ANI Threshold | MashMap2 |  |  |  |  | MashMap3 |  |  |  |  |
| --- | --- | --- | --- | --- | --- | --- | --- | --- | --- | --- | --- |
|  |  | Basepairs mapped (Gbp) | CPU time (m) | Memory (Gb) | ME | MAE | Basepairs mapped (Gbp) | CPU time (m) | Memory (Gb) | ME | MAE |
| Chimpanzee | 95% | 2.80 | 39.76 | **19.95** | -0.25 | 0.29 | **2.81** | **32.76** | 27.07 | **0.01** | **0.22** |
| Chimpanzee | 90% | 2.82 | 118.31 | **24.55** | -0.22 | 0.29 | 2.82 | **51.12** | 36.20 | **0.01** | **0.25** |
| Chimpanzee | 85% | 2.83 | 787.44 | 44.96 | -0.18 | 0.27 | 2.83 | **64.48** | **39.47** | **0.02** | **0.25** |
| Macaque | 95% | 0.38 | 30.0 | **20.83** | **0.29\*** | **0.46** | **1.08** | **28.67** | 28.97 | 0.57* | 0.66 |
| Macaque | 90% | 2.54 | 40.49 | **23.04** | -0.30 | **0.69** | **2.56** | **34.87** | 35.91 | **0.01** | 0.74 |
| Macaque | 85% | 2.60 | 446.71 | **38.13** | -0.24 | **0.74** | **2.61** | **43.74** | 39.49 | **0.05** | 0.87 |

To isolate the effect of the new seeding method, we turned chaining off for both tools. As the Jaccard estimator is known to perform poorly in the presence of many degenerate *k*-mers, results for query regions above and below 80% complexity are reported separately, where complexity is defined as the ratio of observed distinct *k*-mers in a region to *w*. Low-complexity mappings make up for at most 1% and 3% of the mappings for chimpanzee and macaque genomes, respectively. We show the table of the metrics for the low-complexity mappings in Supplementary Table 1.

## 6 Discussion

Minmers are a novel “non-forward” winnowing scheme with a (*w, s*)-window guarantee. Similar to what has been done for other proposed schemes, we have derived formulas (approximate and exact) that describe the scheme’s characteristics. We have replaced minimizers with minmers in MashMap3 and demonstrated that minmers eliminate Jaccard estimator bias and enable new methods to reduce mapping runtime compared to MashMap2. In addition, we show that minmers require substantially less density than minimizers when a (*w, s*)-window guarantee is required.

### The minmer scheme enables sparser sketches

The minimizer winnowing scheme has long been the dominant method for winnowing due to its (*w*, 1)-window guarantee, simplicity, and performance. Other 1-local methods such as strobemers Sahlin (2021) and syncmers Edgar (2021) remove the window guarantee and rely on a random sequence assumption to provide probabilistic bounds on the expected distance between sampled *k*-mers.

Minmers represent a novel class of winnowing schemes that extend the window guarantee of minimizers. Unlike strobemers, syncmers, and other 1-local methods, the minmer scheme guarantees the desired number of *k*-mers will be sampled from every window, so long as it contains at least *s* distinct *k*-mers. This is particularly desirable for accurate Jaccard estimation and the winnowing of low-complexity sequence where the density of sampled *k*-mers from 1-local schemes can vary significantly.

### Minmers yield an unbiased estimator at lower computational costs

Indexing minmers rather than minimizers removes the Jaccard estimator bias present in earlier versions of MashMap. For any window, the set of sampled *k*-mers is guaranteed to be a superset of the bottom-*s* sketch of that window. Therefore, running the minhash algorithm on minmers yields the same estimator as running the minhash algorithm on the full set of *k*-mers.

In addition to the experiments from Belbasi *et al*. (2022), which focus on “ideal” sequences with no repetitive *k*-mers, we also measured the performance of the ANI prediction for different levels of divergence on the human genome across mappings of simulated reads and a sample of mammalian genomes. Our results showed that MashMap3 with minmers not only produced unbiased and more accurate predictions of the ANI than Minimap2 and MashMap2, but it did so in a fraction of the time.

We replicated the behavior of minimizers to under-predict ANI as seen in Belbasi *et al*. (2022) across all experiments. At the same time, in both the simulated reads and empirical genome alignment results, we see that MashMap3 slightly over-predicts the ANI at larger divergences. Further inspection reveals that this is due to indels in the alignment, which are not modeled by the binomial mutation model used to convert the Jaccard to ANI (Supplementary Table 2).

The optimizations to the second stage of mapping combined with the minmer interval indexing leads to significantly better mapping speeds in MashMap3. Relative to Minimap2 and MashMap2, MashMap3 spends a significant amount of time indexing the genome. This, however, serves as an investment for the mapping phase which is significantly faster than MashMap2, particularly at lower ANI thresholds. As an additional feature, MashMap3 provides the option to save the reference index so that users can leverage the increased mappings speeds for previously indexed genomes.

Similar to MashMap2, MashMap3 by default uses the plane-sweep post-processing algorithm described in Jain *et al*. (2018a) to filter out redundant segment mappings. We show that by using the probabilistic filtering method described in Section 4.2.1, we can discard many of these mappings at the beginning of the process as opposed to the end, yielding significant runtime improvements. MashMap3 is significantly more efficient at lower ANI thresholds, which is helpful for detecting more distant homologies. For example, in our human-chimpanzee mapping, we recovered an additional 50 Mbp of mapped sequence by reducing the ANI threshold from 95% to 85% while also completing over 10x quicker than MashMap2. It is also worth noting that the default ANI of MashMap2 and MashMap3 is 85%, and often the ANI of homologies between genomes is not known a priori.

Further motivating the improved efficiency of low ANI thresholds is the fact that thresholds above the true ANI can lead to recovering mappings which over-predict the ANI while discarding those which accurately or under-predict the ANI. This sampling bias leads to an increase in the ANI estimation bias. We see this behavior in the human-macaque alignment with a threshold 95% ANI (Table 2). At lower ANI thresholds, we observe that the majority of mappings are in the 90%-95% ANI range.

### Limitations and future directions

MashMap’s Jaccard-based similarity method tends to overestimate ANI in low-complexity sequences. For downstream alignment applications, the resulting false-positive mappings can be pruned using a chaining or exact alignment algorithm to validate the mappings. Unreliable ANI estimates could also be flagged by using the bottom-*s* sketch to determine the complexity of a segment as described in Cohen and Kaplan (2007), but a sketching method and distance metric that better approximates ANI across all sequence and mutational contexts would be desirable.

An important characteristic of MashMap is the relatively few parameter settings necessary to tune across different use cases. Building on this, we aim to develop a methodology that can find maximal homologies without a pre-determined segment size, similar to the approach of Wang *et al*. (2022b).

## 7 Conclusion

In this work, we proposed and studied the characteristics of the minmer scheme and showed that they belong to the unexplored class of non-forward local schemes, which have the potential to achieve lower densities under the same locality constraints as forward schemes Marçais *et al*. (2018). We derived formulas for the density and approximate spread of minmers, enabling them to be objectively compared to other winnowing schemes.

By construction, minmers, unlike minimizers, enable an unbiased estimation of the Jaccard. We replaced the minimizer winnowing scheme in MashMap2 with minmers and showed that minmers significantly reduce the bias in both simulated and empirical datasets.

Through leveraging the properties of the minmers, we implemented a number of algorithmic improvements in MashMap3. In our experiments, these improvements yielded significantly lower runtimes, particularly in the case when the ANI threshold of MashMap is set to the default of 85%. With the improvements in MashMap3, it is no longer necessary to estimate the ANI of homologies a priori to avoid significantly longer runtimes, making it an ideal candidate for a broad range of comparative genomics applications.

## Acknowledgements

We would like to thank Chirag Jain for helpful discussions and his implementation of the original MashMap software, as well Andrea Guarracino for improvements and discussions. We would also like to thank Nicolae Sapoval and Fritz Sedlazeck for their feedback on the proofs and primate alignments, respectively.

