## Supplemental Material for "Minmers are a generalization of minimizers that enable unbiased local Jaccard estimation"

May 16, 2023

### 1 Supplementary Materials

#### 1.1 Probabilistic filtering for the minhash

We construct a predictor of the numerator of the minhash formula conditioned on the size of the intersection $|\pi_s(A) \cap \pi_s(B_i)|$. This predictor generates a probability distribution for the ANI of a candidate mapping without needing compute the expensive $\pi_s(A \cup B_i)$ step. We start by dividing $\pi_s(A) \cup \pi_s(B_i)$ into two parts where $C_i = \pi_s(A) \cap \pi_s(B_i)$ and $G_i = (\pi_s(A) \cup \pi_s(B_i)) \setminus C_i$ resulting in two sets of size $c_i$ and $2s - c_i$, respectively. The problem can now be formulated as follows: what is the probability that $y$ elements from $C_i$ are also part of the sketch $\pi_s(A \cup B_i)$?

Leveraging the fact that $\pi_s(A \cup B_i) = \pi_s(\pi_s(A) \cup \pi_s(B_i))$ and that all orderings of elements in $\pi_s(A \cup B_i)$ are equally likely, we can view the problem as assigning the $c_i$ shared elements to $2s - c_i$ slots, where the first $s$ slots are considered as a "success" and the remaining $s - c_i$ slots are considered as a "failure" (Supplementary Figure 1).

We have the following formulas:

$$
\begin{aligned}
\Pr(Y_i = y | c_i) &= \text{Hypergeom}_{pdf}(2s - c_i, s, c_i, y) \\
&= \frac{\binom{s}{y}\binom{s-c_i}{c_i-y}}{\binom{2s-c_i}{c_i}} \\
\Pr(Y_i \leq y | c_i) &= \text{Hypergeom}_{cdf}(2s - c_i, s, c_i, y) \\
&= \sum_{i=0}^{y-1} Pr(Y_i = y | c_i)
\end{aligned}
$$

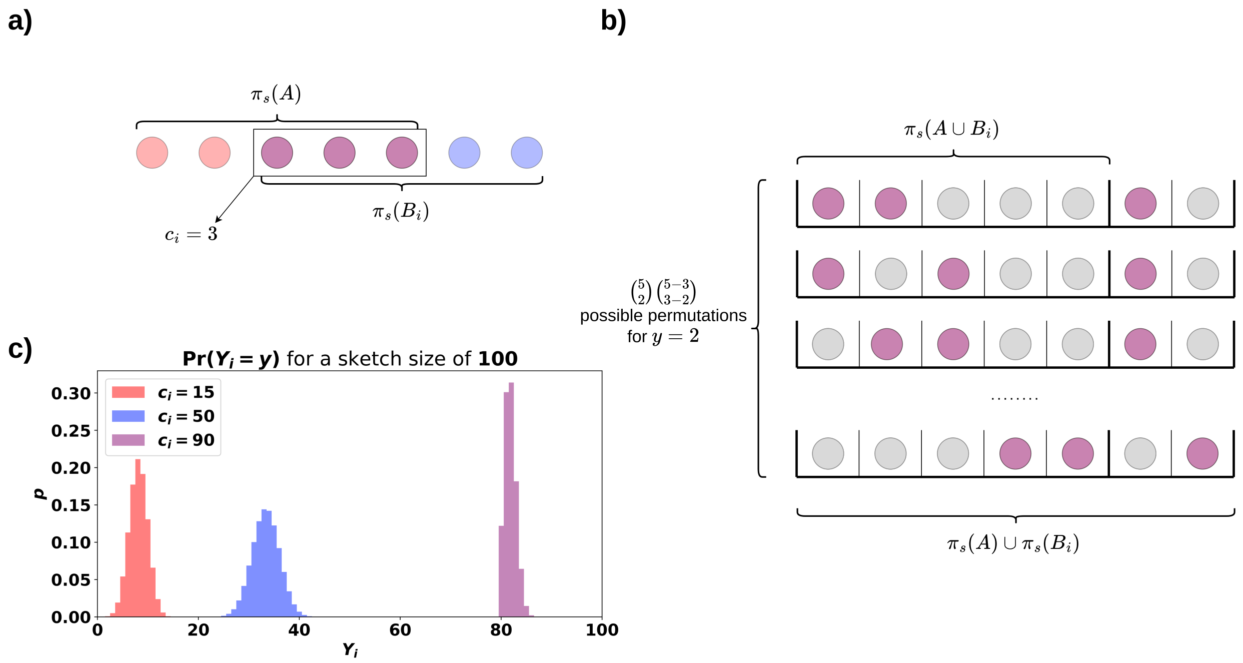


Figure 1: **Estimating the minhash from $\pi_s(A)$ and $\pi_s(B_i)$.** **(a)** Given two sketched sets $\pi_s(A)$ and $\pi_s(B_i)$, we can compute the size of their intersection $c_i$. **(b)** By considering $C_i = \pi_s(A) \cap \pi_s(B_i)$ as purple balls and $G_i = (\pi_s(A) \cap \pi_s(B_i)) \setminus C_i$ as grey balls, we can enumerate all possible permutations of their union such that exactly $y$ purple balls fall within the first $s$ slots. **c)** The distribution of the minhash numerator $Y_i$ for different values of $c_i$ when $s = 100$. The corresponding distribution of the minhash can be obtained by dividing $Y_i$ by the sketch size $s$.

#### 1.2 ANI prediction performance on low-complexity queries

| | | MashMap2 | | | MashMap3 | | |
|---|---|---|---|---|---|---|---|
| Query Species | ANI Threshold | Basepairs mapped (Gbp) | ME | MAE | Basepairs mapped (Gbp) | ME | MAE |
| chimpanzee | 95% | 0.01 | **0.76** | **1.36** | 0.01 | 1.05 | 1.51 |
| chimpanzee | 90% | 0.03 | 4.51 | 4.76 | 0.03 | **4.43** | **4.63** |
| chimpanzee | 85% | 0.04 | 4.85 | 5.11 | 0.04 | **4.81** | **5.03** |
| macaque | 95% | <0.01 | **0.63** | 1.66 | <0.01 | 0.86 | **1.55** |
| macaque | 90% | <0.01 | 2.13 | 2.96 | <0.01 | **0.72** | **1.74** |
| macaque | 85% | 0.05 | 9.79 | 9.88 | **0.08** | **7.98** | **8.03** |

Table 1: **Proportion and accuracy of low-complexity mappings.** MashMap2 and MashMap3 were used to align the human reference genome to chimpanzee and macaque genomes. The number of aligned query query nucleotides from low-complexity segments as well as the ME and MAE of the mappings are reported here.

#### 1.3 Simulated read results and the effects of indels

| Difference Ratio | ONT-95 ME | ONT-98 ME | ONT-99 ME |
|---|---|---|---|
| 20:40:40 | 0.30 | 0.11 | 0.05 |
| 100:00:00 | 0.00 | -0.02 | -0.02 |

Table 2: **The effect of indels on ANI prediction error.** For error rates of 1%, 2%, and 5%, Pbsim was used to generate two datasets, one with a mismatch, insertion, deletion ratio of 20:40:40 and another with mismatches only (100:00:00). ANI was estimated from the Jaccard using the binomial model.

#### 1.4 Minmer density

To obtain the density of the minmer scheme, we inspect how the rank of a $k$-mer changes with each sliding window. In particular, we use the rank of the $k$-mer in its first and last windows, i.e. the windows in which the $k$-mer is just entering and just about to leave. To inspect this, we characterize the distribution of the first rank, the distribution of the final rank given the first rank, and the probability of the rank ever being less than or equal to $s$ given the first and last ranks.

Let $S$ be a sequence of $2w-1$ uniformly random numbers in $[0,1]$. We denote the middle element at position $w$ as $z$, its rank in the leftmost window of size $w$ as $r_1$, and its rank in the rightmost window of size $w$ as $r_w$. Let $C_{r_1,r_w}$ be a conditional indicator r.v. such that $\Pr(C=1|r_1,r_w) = \Pr(C_{r_1,r_w}=1)$ where $C=1$ only if there exists a window of length $w$ in $S$ such that the rank of $z$ in the window is at most $s$. This event corresponds to the element $z$ being a minmer.

**Lemma 1.1.**

$$\Pr(C_{r_1,r_w}) = \begin{cases} \sum_{u=0}^{\delta} \Pr(U=u)\frac{\binom{2u+r_w-r_1}{u+r_w-s}}{\binom{2u+r_w-r_1}{u}} & r_1 > s, r_w > s \\ 1 & otw \end{cases}$$

where $U \sim$ Hypergeometric$(w-1, r_1-1, w-r_w)$ and $\delta = \min(r_1-1, w-r_w)$.

*Proof.* Given the initial rank $r_1$ and the final rank $r_w$, we can model the path of the rank as left and right unit steps on a number line starting at point $r_1$ and ending at $r_w$. At each step in this path, the rank either increases, decreases, or remains the same. The event $C_{r_1,r_w}$ is then equivalent to the event that the path touches the point $s$ on the axis. Let $\omega = \omega_{\text{left}} z \omega_{\text{right}}$ be a sequence of length $2w-1$ representing the elements in $S$. We let $\omega_{left} = ppqpq...$ and $\omega_{\text{right}} = qpqppq...$ where each element is labeled as $p$ if it is less than $z$ and $q$ otherwise. We define $x$ and $y$ as the number of $p$s and $q$s in $\omega_{\text{left}}$, respectively, and similarly $a$ and $b$ are the number of $p$s and $q$s in $\omega_{\text{right}}$, respectively. At step $i$, the rank $z$ can decrease only if $\omega_{\text{left}}[i] = p$ and $\omega_{\text{right}}[i] = q$. Similarly, the rank will increase only if $\omega_{\text{left}}[i] = q$ and $\omega_{\text{right}}[i] = p$. Otherwise, the rank will remain the same. We note that there can be no more than $\max(r_1-1, w-r_w)$ left steps, as $x = r_1-1$ and $b = w-r_w$.

For each of the $x$ $p$s in $\omega_{\text{left}}$, we sample without replacement from $\omega_{\text{right}}$. By considering each sampling of a $q$ as a success, we see that the number of left steps given the initial and final ranks $r_1$ and $r_w$ can be modeled as a hypergeometric random variable $U \sim \text{Hypergeometric}(w-1, x, b)$.

With a set of $u$ left steps, we can calculate the number of right steps $v$ by observing that if we have $u$ $pq$ pairs, then there must be $x-u$ $pp$ pairs, $b-u$ $qq$ pairs, and therefore $y-(b-u) = r_w - r_1 + u$ $qp$ pairs. Given a set of $u$ left steps and $v$ right steps, there are $\binom{u+v}{u}$ total paths. Of these paths, we aim to find the ones which touch point $s$ on the axis. Using the reflection principle Comtet (1974), we observe that there are $\binom{u+v}{u+r_w-s}$ such paths and therefore

$$\frac{\binom{u+v}{u+r_w-s}}{\binom{u+v}{u}} = \frac{\binom{2u+r_w-r_1}{u+r_w-s}}{\binom{2u+r_w-r_1}{u}}$$

□

With the conditional distribution $C_{r_1,r_w}$ at hand, we can define the marginal distribution of $C$.

**Lemma 1.2.**

$$\Pr(C=1) = \frac{1}{w} \sum_{r_1, r_w \in \{1 \ldots w\}^2} \Pr(C=1|r_1, r_w) \Pr(R_w = r_w|r_1)$$

Where $R_1 \sim \text{Uniform}\{1, w\}$ and $R_w|r_1 \sim \text{BetaBinomial}(r_1, w-r_1+1)$ are random variables for the first and last rank of $z$, respectively.

*Proof.* Given $r_1$, the initial rank of $z$, we can use order statistics for uniform distributions to infer that the value of $z$ is sampled from a Beta distributed r.v. $Z \sim \text{Beta}(r_1, w-r_1+1)$. Given the value $z$, we can predict the final rank of $z$ by considering the remaining $w-1$ elements as Bernoulli trials each with probability $z$ of having a lower value than $z$. Therefore, we have that $R_w|z \sim \text{Bin}(w-1, z)$. We can obtain the marginal of $R_w$ via

$$\Pr(R_w = r_w) = \int_0^1 \Pr(R_w = r_w|p) \Pr(Z=z) dp$$

which is the Beta-binomial distribution with $n = w-1$, $\alpha = r_1$ and $\beta = w - r_1 + 1$. □

#### 1.5 Minmer interval density

We will prove the density of minmer intervals in a similar fashion to the proof for minimizers. We define a window of length $w$ as at position $i$ as $W_i$ and say $W_i$ is *charged* if $\pi_s(W_i) \neq \pi_s(W_{i-1})$. Like minimizers, the set of minmers between two adjacent windows can differ by at most one, as only a single minmer can leave the sketch at a time. Unlike minimizers, though, it is possible for a $k$-mer at position $i$ to charge multiple windows by exiting and then re-entering the sketch. Therefore, the number of charged windows in a sequence is at least the number of minmers.

Consider a super-window of $w+1$ $k$-mers starting at position $i-1$ and let $\pi_s(W_i \cup W_{i-1})$ be the lowest $s$ $k$-mers in the super-window. $W_i$ is then not charged if and only if both $x_{i-1} \notin \pi_s(W_i \cup W_{i-1})$ and

$x_{i+w-1} \notin \pi_s(W_i \cup W_{i-1})$. Assuming each position is equally likely to be part of the sketch, the probability of the first and last $k$-mers not being in the sketch is $\binom{w-1}{s}/\binom{w+1}{s}$ and therefore the probability that $W_i$ is charged is

$$\begin{aligned}\Pr(W_i \text{ is charged}) &= 1 - \frac{\binom{w-1}{s}}{\binom{w+1}{s}} \\ &= 1 - \frac{(w-s+1)(w-s)}{w(w+1)}\end{aligned}$$

Assuming independence over windows, we have that the density of charged windows is equal to the probability that any window is charged and therefore the density of minmer intervals is $1 - \frac{(w-s+1)(w-s)}{w(w+1)}$.

#### 1.6 Minmer spread

We now turn our attention to characterizing the distribution of distances between adjacent minmers using a proof described in joriki (2012).

Consider a window of length $w+1$ which contains $s$ sampled $k$-mers and is anchored at the left-most sampled $k$-mer. Assuming a set of $w+1$ unique $k$-mers , we have that each of the $w+1$ $k$-mers is equally likely to be sampled. Let $X_1, ..., X_{s-1}$ be a set of integers randomly sampled from $\{1, ..., w\}$ such that $X_i < X_{i+1}$. We define the distance between $X_i$ and $X_{i+1}$ as $G_i = X_{i+1} - X_i$. We let $X_0 = 0$ represent the first $k$-mer in the window positioned at the first location.

**Lemma 1.3.** $\Pr(G_i = d) = \frac{\binom{w-d}{s-2}}{\binom{w}{s-1}}$

*Proof.* Let us consider our $w+1$ unique sorted integers arranged on a circle instead of a line. We then "cut" the circle at any one of the $s$ sampled integers and renumber the $w$ remaining integers starting from 1 after the cut. There are now $s-1$ integers uniformly sampled from $\{1, ..., w\}$. By fixing the first sample at position $d$ and enforcing that all $s-2$ remaining integers are sampled from $\{d+1, ..., w\}$, we see that there are $\binom{w-d}{s-2}$ such samples. Given that there are $\binom{w}{s-1}$ ways to sample the $s-1$ integers, the distance $d$ between the cut and the first sampled point is then distributed as $\frac{\binom{w-d}{s-2}}{\binom{w}{s-1}}$. As this analysis is symmetric for any "cut," we claim that the distribution of all $G_i$ are identical. □

While the analysis above is conditioned on the case where we have $s$ uniformly random chosen positions, the number of sampled positions varies across windows and is only lower-bounded by $s$. If we replace $s$ with the expected number of minmers in the window, $d_f$, we can obtain an approximation of the distribution of distances (Figure 2). A more rigorous analysis, which is beyond the scope of this work, would require a distribution for the number of sampled positions in a window rather than just the expectation.

Unfortunately, this distribution is not that useful on its own. Given that the distribution of the distance is the same across all points, we have that $(s+1)\,\mathbb{E}[G_i] = w+1$ and therefore $\mathbb{E}[G_i] = (w+1)/(s+1)$. Even more interesting than the expectation, though, are the order statistics of $G_i$, such as $\max G_i$.

In *Order Statistics* David and Nagaraja (2004), a similar problem is studied where a rope of length 1 is cut at $n$ randomly selected locations. The authors show that the expected length of the longest segment is $H_{n+1}/(n+1)$, where $H_n$ is the $n$th harmonic number. The details of the problem we describe above are slightly different, as the “cut-points” are selected from a set of integers without replacement as opposed to sampled from $[0,1]$. We can use this to define $\bar{\mathcal{G}}_i$, an estimator for $\max G_i$,

$$\bar{\mathcal{G}}_i = (w+1)\frac{H_{d_f+1}}{d_f+1}$$

As $w$ grows, the effect of sampling without replacement grows smaller and the error of $\bar{\mathcal{G}}_i$ becomes solely from the fact that $d_f$ is only an expectation of the number of minmers in a window.

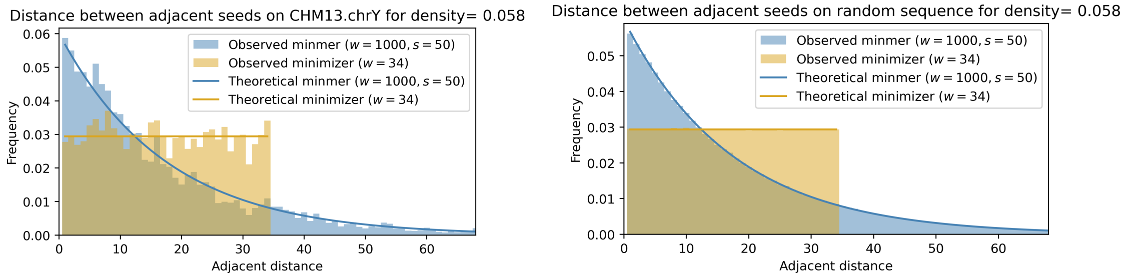


(a) (b)

Figure 2: The spread of minmers and minimizers under similar densities on the human Y-chromosome (a) and a simulated random sequence (b).
